# AlphaFold2 and AlphaFold3 leads to significantly different results in human-parasite interaction prediction

**DOI:** 10.1101/2024.09.19.613643

**Authors:** Burcu Ozden, Buse Şahin, Ezgi Karaca

## Abstract

**Motivation:** Parasitic diseases pose a significant global health challenge with substantial socioeconomic impact, especially in developing countries (Dampier et al., 2009; Mehmood et al., 2017; Merrifield et al., 2016). Combatting these diseases is difficult due to two factors: the resistance developed by parasites to existing drugs and the underfunded research efforts to resolve host-parasite interactions (Cuesta-Astroz & Oliveira, 2018). To address the latter, we focused on 276 human-parasite domain-domain interaction predictions, involving 15 parasitic species known to cause infections in humans (Table S1, Cuesta-Astroz et al., 2019). We aimed to validate these interactions by using all AlphaFold2-Multimer (AF2) and AlphaFold3 (AF3) (Jumper et al., 2021; Abramson et al., 2024). Our initial hypothesis was that the interactions with an AF confidence score ≥ 0.8 could serve as high confident templates for further research. However, through systematic modeling of these interactions, we observed a striking discrepancy: AF3 confidence assignment behaves significantly differently from that of AF2. In this study, we investigate the underlying causes of this difference.

## Materials and Methods

### Dataset Preparation

This study is based on a human–parasite protein–protein interaction set, predicted for 15 distinct eukaryotic parasites (Table S1, Cuesta-Astroz et al., 2019). The species included in this work are *T. spiralis, T. gondii, T. brucei, S. mansoni, P. vivax, P. knowlesi, L. infantum, L. donovani, L. braziliensis, C. parvum, G. lamblia, T. cruzi, P. falciparum, C. hominis*, and *L. mexicana*. The original data sset consisted of 280 domain–domain interactions. From this set, four pairs were excluded due to their combined protein sequence lengths exceeding 1,500 residues. The domain-based fasta files of the final of 276 interaction set are provided at https://aperta.ulakbim.gov.tr/record/263582.

### Structure Prediction

We run AF 2.1, 2.2, and 2.3 versions through the MassiveFold pipeline (Raouraoua *et al*., 2024). The Multiple Sequence Alignment (MSA) generation was performed with MMseqs2 over the full database preset (UniRef90, UniRef30_2021_03, UniProt, BFD, MGnify). The relaxation step was disabled. The run parameters are provided in Supplementary Table 2. AF3 predictions were performed by using the official AlphaFold Server (Abramson et al., 2024) with default settings. For all runs, we used a single seed.

To ensure a consistent comparison across AF versions, the confidence scores for all models were standardized using [0.8×ipTM + 0.2×pTM] confidence formula. All the scores and are provided as a csv file together with this preprint (*all_scores*.*csv*).

### Analysis Tools

DockQ was used to assess model accuracy (Basu and Wallner, 2016). Model comparisons were classified according to CAPRI standards: DockQ ≥ 0.80: high quality; DockQ ≥ 0.49: medium quality; DockQ ≥ 0.23: acceptable quality. The acquired DockQ values are provided as a csv file together with this preprint (*dockq_scores*.*csv*).

The effective sequence count (eff_nseq) values were calculated with MMseqs2. For each complex, we calculated a complex-specific geometric Neff, as described in CASP15 Assembly Assessment paper (Ozden et al. 2025).

Data analysis and visualizations were performed by using Python (pandas, NumPy, matplotlib, seaborn). Structural representations were prepared using PyMOL v3.1.3.1.

## RESULTS AND DISCUSSION

### Across All AF Versions AF3 Produces the Lowest Confidence Scores

First, to assess inter-version confidence score distributions, we modeled 276 host-parasite domain-domain interactions using all available AlphaFold-Multimer (AF) versions, i.e. AF2.1, AF2.2, AF2.3, and AF3. Comparison of the top-ranking prediction scores (Figure 1) revealed that AF3 consistently produced the lowest confidence, with a median of 0.28, significantly lower than AF2.1 (0.45), AF2.2 (0.39), and AF2.3 (0.36). Given the bioinformatically predicted nature of the initial interaction dataset, we applied a conservative confidence threshold of ≥0.8 to select high-confidence cases for detailed analysis. This threshold isolated 65 unique interaction pairs where at least one model achieved the threshold, with the model counts being: AF2.1 (57), AF2.3 (41), AF2.2 (39), and AF3 (14) (Figure 1). The subsequent sections focus on a detailed analysis of these selected “confident” models.

**Figure 1.**
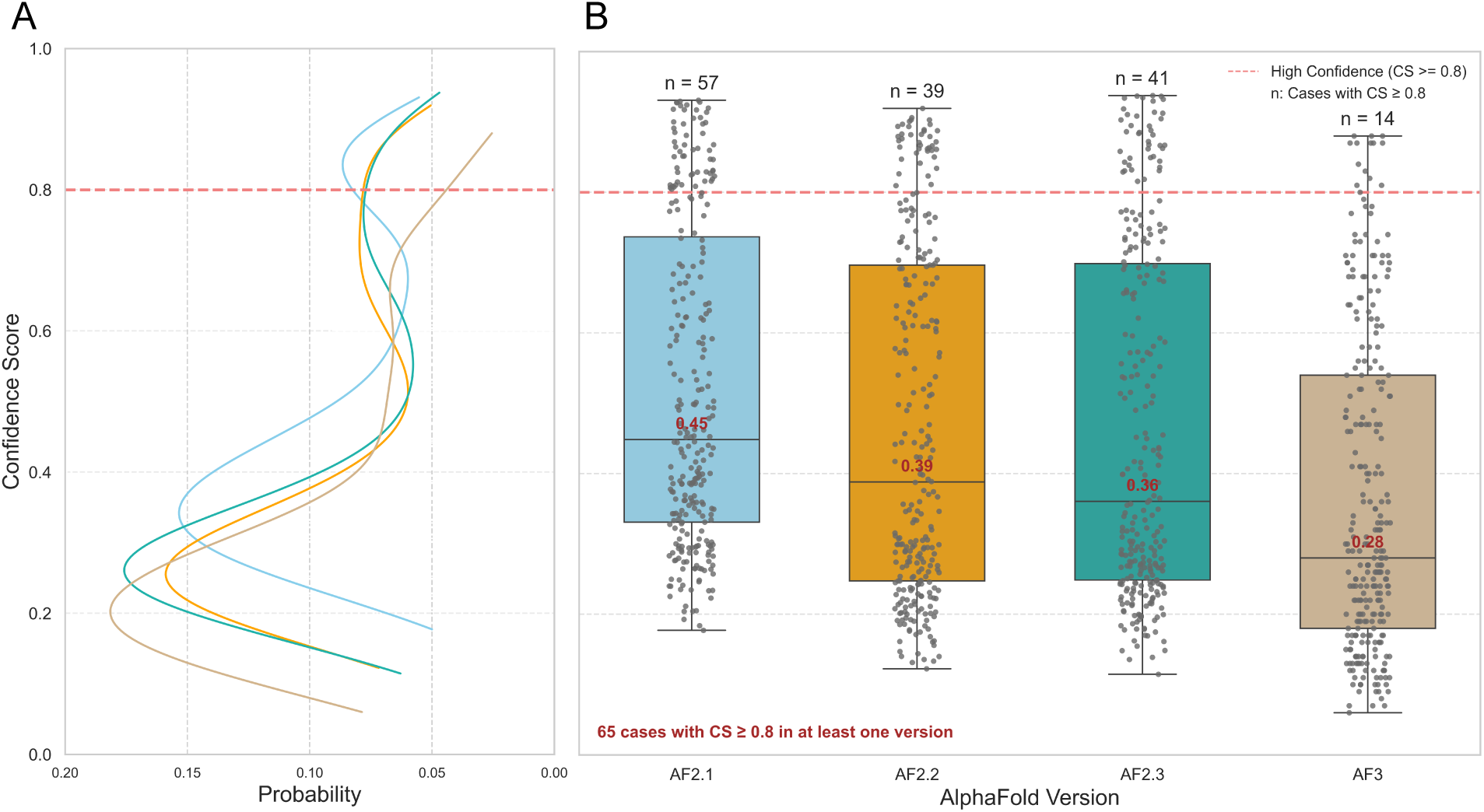
**(A)** Probability and **(B)** Box-and-whisker plot distributions of confidence scores (top-ranked model), generated by each AF version.

### AF3 Confidence Scores Reflect Lower Correlation with the AF2 Ones

To investigate the reason for the differential AF3 scoring, we generated pairwise scatter plots of confidence scores for all AF versions. This analysis revealed a high degree of agreement among AF2 versions. For example, the top confidence scores generated by AF2.2 and AF2.3 reflected an R^2^ of 0.83 (Figure 2A). In contrast, comparisons involving AF3 showed substantially lower correlations with all AF2 versions, peaking at an R^2^ of 0.54 (AF3 vs. AF2.3, Figure 2B).

**Figure 2.**
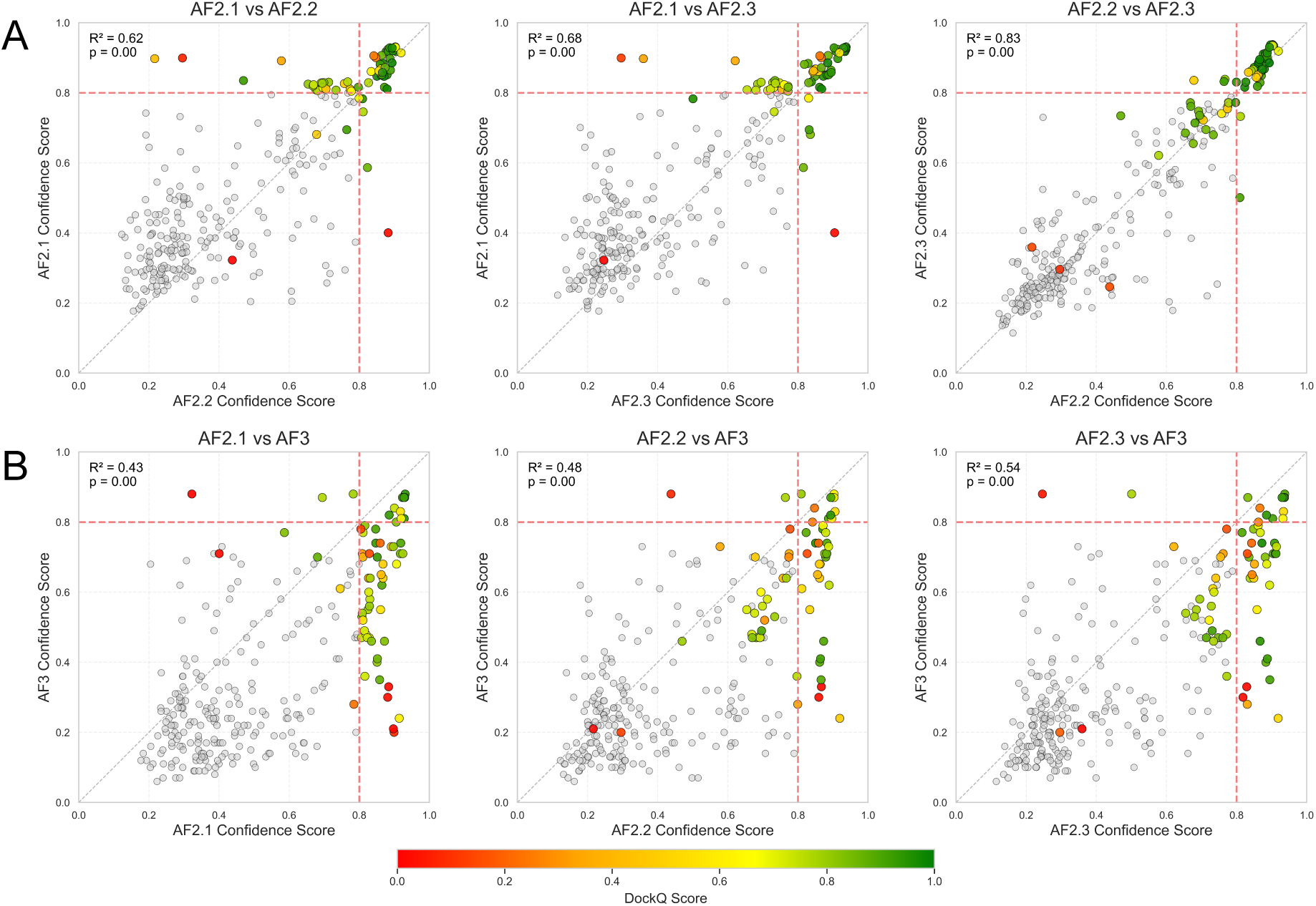
Confidence score comparisons A. AF2 vs. AF2, and B. AF2 and AF3 versions. The “confident” 65 cases are colored according to the DockQ score of the compared models.

We then calculated the structural similarities for the 65 high-confidence models using DockQ scores and mapped these onto the scatter plots (Figure 2). We anticipated that pair with dissimilar confidence scores would yield models with low structural similarities. However, in the AF3 versus AF2 comparisons, we identified several *outliers*, where significantly different confidence scores (e.g., AF3 conf-score < 0.8 & AF2 conf-score > 0.8) produced structures with high DockQ scores.

Figure 3 presents two exemplary outlier cases, where AF2 and AF3 predicts similar structures with very different confidence scores. This scoring divergence is visually confirmed by the pLDDT coloring: AF2 models display high confidence, whereas AF3 predictions feature several cyan and yellow (lower-confidence) regions across the interaction partners. In the following section, we investigate the underlying cause for this scoring discrepancy.

**Figure 3.**
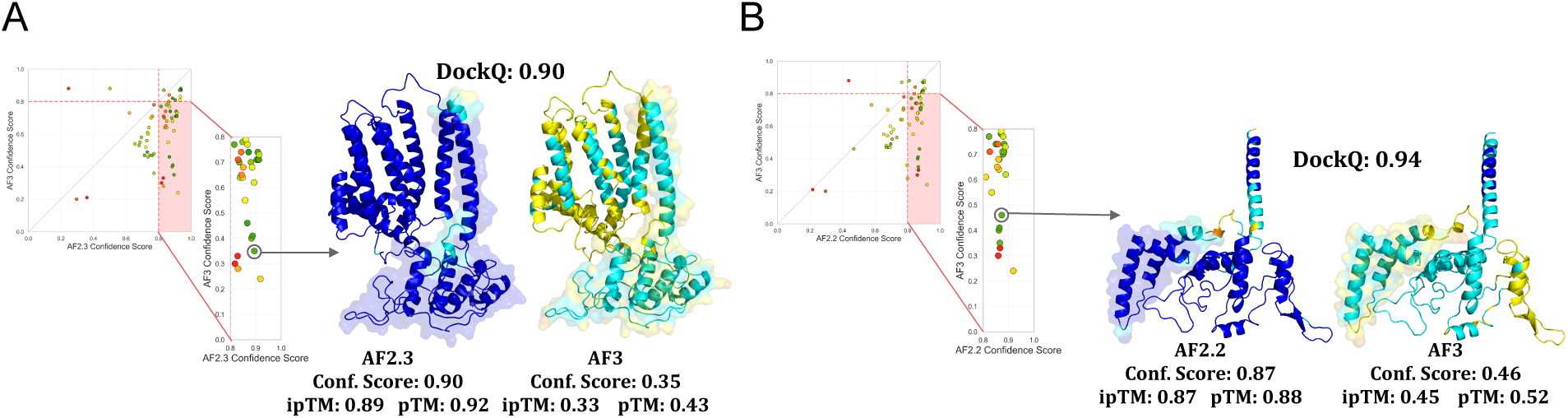
Structural comparison of two outlier cases: **A.** The ENSP00000354554_TGME49_046540 complex and **B**. ENSP00000309565_TGME49_046540 complex from the human-T. gondii host-pathogen interaction set. Complexes are colored according to their plDDT scores. In each complex, one of the monomers is represented as transparent surface.

### Exploring the Reason Behind the AF2/AF3 Scoring Discrepancy

To understand the scoring discrepancy between AF2 and AF3, we first analyzed the confidence score distributions per parasitic species (Figure 4). This analysis showed that for all species where at least one structure surpassing the 0.8 confidence threshold, AF3 produced smaller confidence scores. Interestingly, for a few host–parasite pairs, such as human–*C. hominis*, human–*T. brucei*, and human–*L. Donovani*, most of the interactions were predicted to be unreliable by all AF versions (Supplementary Table 3). This outcome highlights that there is no species dependency on the scoring results we observed.

**Figure 4.**
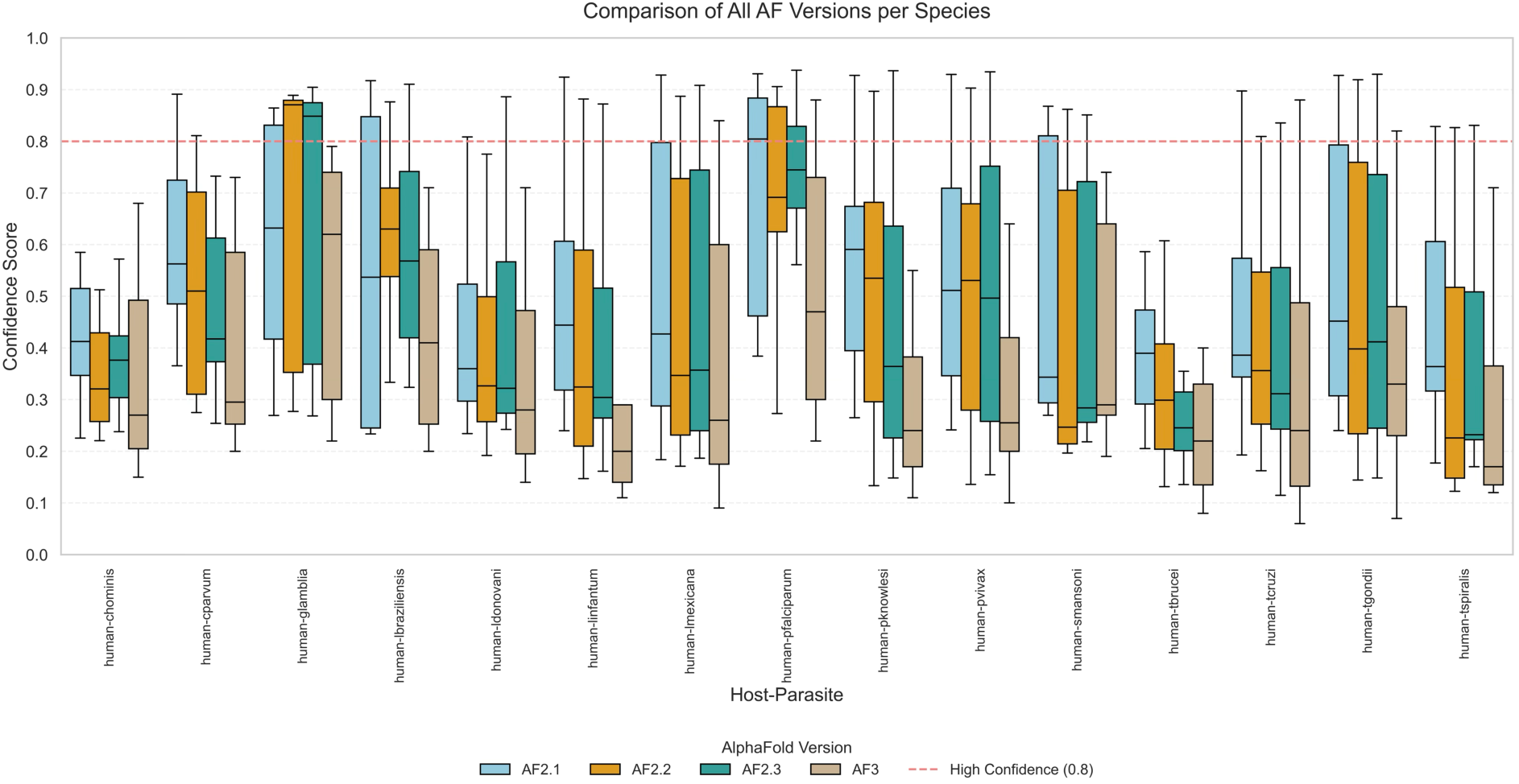
Box-and-whisker plot distributions of confidence scores (top-ranked model) across different species generated by each AF version (AF2-M v1, v2, v3, and AF3). The red dashed line at 0.8 indicates the threshold for high-confidence predictions.

We further investigated the relationship between the MSA depth, and the confidence scores generated by different AF versions. To quantify the MSA content for each dimeric pair, we calculated the geometric mean of the effective number of sequences Neff. We then binned the confidence scores (Figure 5) and analyzed whether a correlation exists between the produced confidence scores and the MSA depth. Surprisingly, this analysis demonstrated that the high-confidence bin ([0.8-1.0]) is populated by cases with MSA depths as shallow as those found in the least-confidence bin ([0.0-0.2]). This indicates that the observed confidence score discrepancy is not a function of the MSA content.

**Figure 5.**
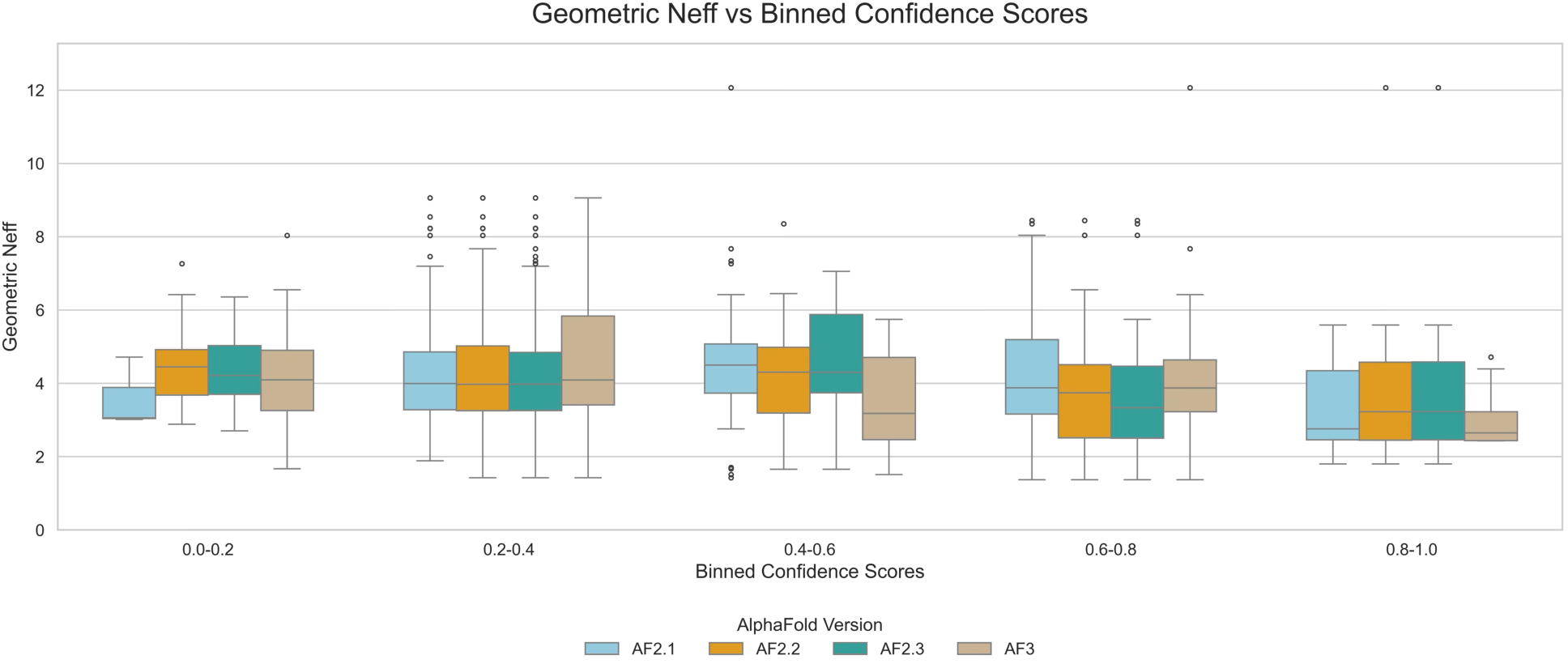
Grouped box-and-whisker plots showing the relationship between Geometric Neff (effective sequence count) and binned confidence scores across AF versions.

As the next step, we performed full-length sequence predictions for the two problematic cases presented in Figure 3. Interestingly, when the full-length sequence context was introduced, we observed a significant confidence boost in AF3 (Figure 6), whereas AF2 scoring remained unchanged. More critically, in both cases, the predicted domain interaction interfaces remained the same despite the score increase. This finding provides the first solid hint towards understanding the reason behind AF3’s differential behavior in scoring: AF3’s scoring is sensitive to the broader sequence context outside of the predicted domain boundaries. However, our efforts in generalizing this outcome across the entire dataset did not result in a consistent observation (data not shown), indicating that it is difficult to predict what AF3 considers to be the *complete* for its confidence calculation.

**Figure 6.**
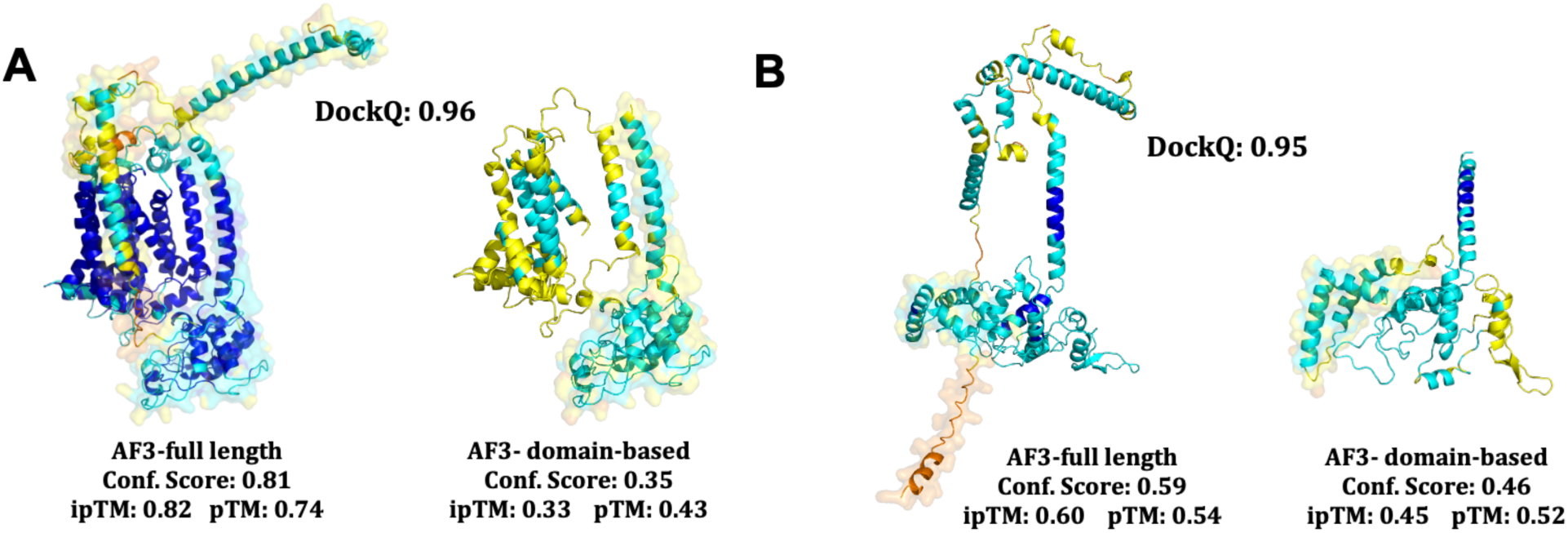
Full-length structural comparison of non-consistently predicted models. Comparison of A. ENSP00000354554_TGME49_046540 complex and B. ENSP00000309565_TGME49_046540 complex from the human-T. gondii host-pathogen pairs. Complexes are colored according to their residue plDDT scores.

## Conclusion

In the end, the use of a host-parasite interaction prediction set led us to discover that AF3 scoring changes significantly with the broader sequence context. On one hand, this behavior is understandable since AF3 is trained to model non-protein systems (including nuclei acids, ligands, and ions). On the other hand, since our experiments showed that it is difficult to generalize what AF3 considers complete, its confidence score can become unpredictable when applied to unknown or unexplored biological systems, like ours. We, therefore, conclude that for the validation of specific protein-protein interaction predictions, such as the focused domain-domain assessments performed here, it would be safer to use AF2.

## Supporting information

dockq_scores

all_scores

## Acknowledgements

The dataset is adapted from our previous TROPIC project (Project number 120N799), funded by Turkish and Colombian Scientific Research Councils (TÜBİTAK and MINCIENCIAS). AF2 models were generated on MareNostrum5 by using MassiveFold (Raouraoua et al.,2025) and AF3 models were generated on AlphaFold webserver (https://alphafoldserver.com/). We wholeheartedly thank the AlphaFold team for their kind help in understanding AF3’s behavior in scoring.

## Notes

### Competing Interest Statement

The authors have declared no competing interest.

### Summary of Updates

The figures are updated, as some of them got corrupted during the upload process.

