## Supporting Information for "AlphaFold2 and AlphaFold3 leads to significantly different results in human-parasite interaction prediction"

**Table S1. Uniprot IDs of Human-Parasite Interaction Predictions.**

| No | Parasite name | Host id | Parasite id | No | Parasite name | Host id | Parasite id |
| --- | --- | --- | --- | --- | --- | --- | --- |
| 1 | *T.spiralis* | ENSP00000258091 | EFV60556 | 139 | *P.knowlesi* | ENSP00000287022 | PKH_124300 |
| 2 | *T.spiralis* | ENSP00000346088 | EFV55619 | 140 | *P.knowlesi* | ENSP00000304697 | PKH_123930 |
| 3 | *T.spiralis* | ENSP00000396937 | EFV54008 | 141 | *P.knowlesi* | ENSP00000389103 | PKH_041010 |
| 4 | *T.spiralis* | ENSP00000233893 | EFV60556 | 142 | *P.knowlesi* | ENSP00000223641 | PKH_041010 |
| 5 | *T.spiralis* | ENSP00000215071 | EFV54008 | 143 | *P.knowlesi* | ENSP00000277865 | PKH_131960 |
| 6 | *T.spiralis* | ENSP00000393393 | EFV55619 | 144 | *P.knowlesi* | ENSP00000262061 | PKH_040280 |
| 7 | *T.spiralis* | ENSP00000322419 | EFV55619 | 145 | *P.knowlesi* | ENSP00000317159 | PKH_124300 |
| 8 | *T.gondii* | ENSP00000279022 | TGME49_001130 | 146 | *P.knowlesi* | ENSP00000358126 | PKH_040940 |
| 9 | *T.gondii* | ENSP00000279022 | TGME49_001130 | 147 | *P.knowlesi* | ENSP00000263354 | PKH_040940 |
| 10 | *T.gondii* | ENSP00000279022 | TGME49_001130 | 148 | *P.knowlesi* | ENSP00000354554 | PKH_124300 |
| 11 | *T.gondii* | ENSP00000279022 | TGME49_001130 | 149 | *P.knowlesi* | ENSP00000294179 | PKH_040940 |
| 12 | *T.gondii* | ENSP00000279022 | TGME49_001130 | 150 | *P.knowlesi* | ENSP00000054666 | PKH_040940 |
| 13 | *T.gondii* | ENSP00000279022 | TGME49_001130 | 151 | *P.knowlesi* | ENSP00000249647 | PKH_040940 |
| 14 | *T.gondii* | ENSP00000279022 | TGME49_001130 | 152 | *P.knowlesi* | ENSP00000227378 | PKH_071520 |
| 15 | *T.gondii* | ENSP00000279022 | TGME49_001130 | 153 | *P.knowlesi* | ENSP00000249356 | PKH_071520 |
| 16 | *T.gondii* | ENSP00000279022 | TGME49_001130 | 154 | *P.knowlesi* | ENSP00000215730 | PKH_040940 |
| 17 | *T.gondii* | ENSP00000279022 | TGME49_001130 | 155 | *P.knowlesi* | ENSP00000259469 | PKH_041010 |
| 18 | *T.gondii* | ENSP00000279022 | TGME49_001130 | 156 | *P.knowlesi* | ENSP00000261558 | PKH_040280 |
| 19 | *T.gondii* | ENSP00000279022 | TGME49_001130 | 157 | *P.falciparum* | ENSP00000249356 | PFI0875w |
| 20 | *T.gondii* | ENSP00000279022 | TGME49_001130 | 158 | *P.falciparum* | ENSP00000297185 | PFI0875w |
| 21 | *T.gondii* | ENSP00000279022 | TGME49_001130 | 159 | *P.falciparum* | ENSP00000302961 | PFI0875w |
| 22 | *T.gondii* | ENSP00000279022 | TGME49_001130 | 160 | *P.falciparum* | ENSP00000277865-PF02812 | PF14_0286 |
| 23 | *T.gondii* | ENSP00000279022 | TGME49_001130 | 161 | *P.falciparum* | ENSP00000265028 | PFI0875w |
| 24 | *T.gondii* | ENSP00000279022 | TGME49_001130 | 162 | *P.falciparum* | ENSP00000277865-PF00208 | PF14_0286 |
| 25 | *T.gondii* | ENSP00000279022 | TGME49_001130 | 163 | *P.falciparum* | ENSP00000310219 | PFI0875w |
| 26 | *T.gondii* | ENSP00000279022 | TGME49_001130 | 164 | *P.falciparum* | ENSP00000227378 | PFI0875w |
| 27 | *T.gondii* | ENSP00000279022 | TGME49_001130 | 165 | *P.falciparum* | ENSP00000366179 | PFI0875w |
| 28 | *T.gondii* | ENSP00000279022 | TGME49_001130 | 166 | *P.falciparum* | ENSP00000365991 | PFI0875w |
| 29 | *T.gondii* | ENSP00000279022 | TGME49_001130 | 167 | *P.falciparum* | ENSP00000344818 | PFC0510w |
| 30 | *T.gondii* | ENSP00000279022 | TGME49_001130 | 168 | *P.falciparum* | ENSP00000327589-PF00208 | PF14_0286 |
| 31 | *T.gondii* | ENSP00000279022 | TGME49_001130 | 169 | *P.falciparum* | ENSP00000364802 | PFI0875w |
| 32 | *T.gondii* | ENSP00000279022 | TGME49_001130 | 170 | *P.falciparum* | ENSP00000362329 | PFC0710w.2 |
| 33 | *T.cruzi* | ENSP00000215956 | XP_815664.1-PF00467 | 171 | *P.falciparum* | ENSP00000343885 | PFC0710w.2 |
| 34 | *T.cruzi* | ENSP00000215956 | XP_815664.1-PF01777 | 172 | *P.falciparum* | ENSP00000357998 | PFI0875w |
| 35 | *T.cruzi* | ENSP00000253788-PF00467 | XP_815664.1-PF01777 | 173 | *P.falciparum* | ENSP00000318687 | PFI0875w |
| 36 | *T.cruzi* | ENSP00000253788-PF01777 | XP_815664.1-PF00467 | 174 | *P.falciparum* | ENSP00000327589-PF02812 | PF14_0286 |
| 37 | *T.cruzi* | ENSP00000265100 | XP_815664.1-PF00467 | 175 | *P.falciparum* | ENSP00000388107 | PFC0510w |
| 38 | *T.cruzi* | ENSP00000272418 | XP_803946.1 | 176 | *P.falciparum* | ENSP00000384144 | PFI0875w |
| 39 | *T.cruzi* | ENSP00000293842 | XP_815664.1-PF00467 | 177 | *P.falciparum* | ENSP00000367623 | PFI0875w |
| 40 | *T.cruzi* | ENSP00000293842 | XP_815664.1-PF01777 | 178 | *L.mexicana* | ENSP00000244007 | XP_003872412.1-PF00454 |
| 41 | *T.cruzi* | ENSP00000296581 | XP_808372.1 | 179 | *L.mexicana* | ENSP00000272317 | XP_003876321.1 |
| 42 | *T.cruzi* | ENSP00000309830 | XP_815664.1-PF00467 | 180 | *L.mexicana* | ENSP00000304697 | XP_003876321.1 |
| 43 | *T.cruzi* | ENSP00000318646 | XP_803946.1 | 181 | *L.mexicana* | ENSP00000335153-PF02518 | XP_003872486.1-PF13589 |
| 44 | *T.cruzi* | ENSP00000319341 | XP_808372.1 | 182 | *L.mexicana* | ENSP00000346037 | XP_003876321.1 |
| 45 | *T.cruzi* | ENSP00000355315 | XP_815664.1_PF00467 | 183 | *L.mexicana* | ENSP00000352336 | XP_003872412.1-PF00454 |
| 46 | *T.cruzi* | ENSP00000361076 | XP_815664.1_PF00467 | 184 | *L.mexicana* | ENSP0000032587-PF02518 | XP_003872486.1-PF13589 |
| 47 | *T.cruzi* | ENSP00000378160 | XP_815664.1-PF01777 | 185 | *L.mexicana* | ENSP00000310219 | XP_003876961.1 |
| 48 | *T.cruzi* | ENSP00000380349 | XP_816718.1 | 186 | *L.mexicana* | ENSP00000322419 | XP_003876321.1 |
| 49 | *T.cruzi* | ENSP00000388107-PF01020 | XP_815664.1 | 187 | *L.mexicana* | ENSP00000246957_1 | XP_003872486.1-PF13589 |
| 50 | *T.cruzi* | ENSP00000412566 | XP_808372.1 | 188 | *L.mexicana* | ENSP00000264864 | XP_003872412.1-PF00613 |
| 51 | *T.cruzi* | ENSP00000419449 | XP_803946.1 | 189 | *L.mexicana* | ENSP00000244007 | XP_003872412.1-PF00613 |
| 52 | *T.cruzi* | ENSP00000433919 | XP_808372.1 | 190 | *L.mexicana* | ENSP00000264864 | XP_003872412.1 |
| 53 | *T.cruzi* | ENSP00000472998 | XP_808372.1 | 191 | *L.mexicana* | ENSP00000246957_1-PF00183 | XP_003872486.1-PF00183 |
| 54 | *T.cruzi* | ENSP00000310596 | XP_808372.1 | 192 | *L.mexicana* | ENSP00000325875-PF02518 | XP_003872486.1-PF00183 |
| 55 | *T.cruzi* | ENSP00000301838 | XP_816718.1 | 193 | *L.mexicana* | ENSP00000297185 | XP_003876961.1 |
| 56 | *T.cruzi* | ENSP00000306223 | XP_808372.1 | 194 | *L.mexicana* | ENSP00000302961 | XP_003876961.1 |
| 57 | *T.cruzi* | ENSP00000291295 | XP_810491.1 | 195 | *L.mexicana* | ENSP00000325875-PF00183 | XP_003872486.1-PF00183 |
| 58 | *T.cruzi* | ENSP00000230050 | XP_815664.1-PF01777 | 196 | *L.mexicana* | ENSP00000227378 | XP_003876961.1 |
| 59 | *T.cruzi* | ENSP00000378160 | XP_815664.1-PF00467 | 197 | *L.mexicana* | ENSP00000335153-PF00183 | XP_003872486.1-PF00183 |
| 60 | *T.cruzi* | ENSP00000302896 | XP_815664.1 | 198 | *L.mexicana* | ENSP00000335153-PF02518 | XP_003872486.1-PF00183 |
| 61 | *T.cruzi* | ENSP00000234310 | XP_810491.1 | 199 | *L.mexicana* | ENSP00000335304-PF00198 | XP_003874617.1-PF00198 |
| 62 | *T.cruzi* | ENSP00000300413 | XP_808372.1 | 200 | *L.mexicana* | ENSP00000335304-PF00364 | XP_003874617.1-PF00364 |
| 63 | *T.cruzi* | ENSP00000230050 | XP_815664.1-PF00467 | 201 | *L.mexicana* | ENSP00000357998 | XP_003876961.1 |
| 64 | *T.cruzi* | ENSP00000274606 | XP_815664.1-PF01777 | 202 | *L.mexicana* | ENSP00000359665 | XP_003872412.1-PF00613 |
| 65 | *T.cruzi* | ENSP00000272298 | XP_810491.1 | 203 | *L.mexicana* | ENSP00000346001 | XP_003876321.1 |
| 66 | *T.cruzi* | ENSP00000265100 | XP_815664.1-PF01777 | 204 | *L.mexicana* | ENSP00000339795 | XP_003876321.1-PF08079 |
| 67 | *T.cruzi* | ENSP00000252543 | XP_815664.1 | 205 | *L.mexicana* | ENSP00000352336 | XP_003872412.1-PF00613 |
| 68 | *T.cruzi* | ENSP00000233946 | XP_816718.1 | 206 | *L.mexicana* | ENSP00000344818 | XP_003876321.1 |
| 69 | *T.cruzi* | ENSP00000274606 | XP_815664.1-PF00467 | 207 | *L.mexicana* | ENSP00000359665 | XP_003872412.1 |
| 70 | *T.cruzi* | ENSP00000337459 | XP_810491.1 | 208 | *L.mexicana* | ENSP00000364802 | XP_003876961.1 |
| 71 | *T.cruzi* | ENSP00000363018 | XP_815664.1-PF01777 | 209 | *L.mexicana* | ENSP00000365991 | XP_003876961.1 |
| 72 | *T.cruzi* | ENSP00000369689 | XP_810491.1 | 210 | *L.mexicana* | ENSP00000375730 | XP_003876321.1-PF08079 |
| 73 | *T.cruzi* | ENSP00000341885-PF03719 | XP_803946.1 | 211 | *L.mexicana* | ENSP00000369134 | XP_003874617.1-PF00198 |
| 74 | *T.cruzi* | ENSP00000460871 | XP_815664.1-PF01777 | 212 | *L.mexicana* | ENSP00000384144 | XP_003876961.1 |
| 75 | *T.cruzi* | ENSP00000345957 | XP_803946.1 | 213 | *L.infantum* | ENSP00000216727 | XP_001469711.1 |
| 76 | *T.cruzi* | ENSP00000342374 | XP_808372.1 | 214 | *L.infantum* | ENSP00000309334 | XP_001463268.1 |
| 77 | *T.cruzi* | ENSP00000363018 | XP_815664.1-PF00467 | 215 | *L.infantum* | ENSP00000360525 | XP_001469711.1 |
| 78 | *T.cruzi* | ENSP00000361076 | XP_815664.1-PF01777 | 216 | *L.infantum* | ENSP00000221801 | XP_001463268.1 |
| 79 | *T.cruzi* | ENSP00000315299 | XP_810491.1 | 217 | *L.infantum* | ENSP00000196551 | XP_001463268.1 |
| 80 | *T.cruzi* | ENSP00000345156 | XP_815664.1-PF00467 | 218 | *L.infantum* | ENSP00000249356 | XP_001470161.1 |
| 81 | *T.cruzi* | ENSP00000341885-PF00333 | XP_803946.1 | 219 | *L.infantum* | ENSP00000227378 | XP_001470161.1 |
| 82 | *T.cruzi* | ENSP00000349467 | XP_810491.1 | 220 | *L.infantum* | ENSP00000262225 | XP_001469004.1 |
| 83 | *T.cruzi* | ENSP00000356156 | XP_810491.1 | 221 | *L.infantum* | ENSP00000252543 | XP_001463268.1 |
| 84 | *T.cruzi* | ENSP00000412324 | XP_810491.1 | 222 | *L.infantum* | ENSP00000260443 | XP_001463268.1 |
| 85 | *T.cruzi* | ENSP00000435777 | XP_803946.1 | 223 | *L.infantum* | ENSP00000300291 | XP_001469711.1 |
| 86 | *T.cruzi* | ENSP00000460871 | XP_815664.1-PF00467 | 224 | *L.infantum* | ENSP00000265100 | XP_001463268.1 |
| 87 | *T.brucei* | ENSP00000311430 | EAN76659 | 225 | *L.infantum* | ENSP00000339095 | XP_001463268.1 |
| 88 | *T.brucei* | ENSP00000341885 | EAN76659 | 226 | *L.infantum* | ENSP00000341730 | XP_001463268.1 |
| 89 | *T.brucei* | ENSP00000421280 | EAN78692 | 227 | *L.infantum* | ENSP00000335321 | XP_001469711.1 |
| 90 | *T.brucei* | ENSP00000389103 | AAZ10133 | 228 | *L.infantum* | ENSP00000345156 | XP_001463268.1 |
| 91 | *T.brucei* | ENSP00000256001 | EAN78692 | 229 | *L.infantum* | ENSP00000346015 | XP_001463268.1 |
| 92 | *T.brucei* | ENSP00000258052 | AAZ11482 | 230 | *L.infantum* | ENSP00000363018 | XP_001463268.1 |
| 93 | *T.brucei* | ENSP00000380427 | EAN78692 | 231 | *L.infantum* | ENSP00000378160 | XP_001463268.1 |
| 94 | *T.brucei* | ENSP00000339095 | EAN76659 | 232 | *L.infantum* | ENSP00000361626 | XP_001469711.1 |
| 95 | *T.brucei* | ENSP00000360034 | EAN76659 | 233 | *L.infantum* | ENSP00000418082 | XP_001463268.1 |
| 96 | *T.brucei* | ENSP00000362744 | EAN76659 | 234 | *L.donovani* | ENSP00000313199 | XP_003865434.1 |
| 97 | *T.brucei* | ENSP00000252725 | EAN78692 | 235 | *L.donovani* | ENSP00000360525 | XP_003865434.1 |
| 98 | *T.brucei* | ENSP00000346067 | EAN76659 | 236 | *L.donovani* | ENSP00000361626 | XP_003865434.1 |
| 99 | *T.brucei* | ENSP00000352918 | EAN78692 | 237 | *L.donovani* | ENSP00000292123 | XP_003862748.1 |
| 100 | *T.brucei* | ENSP00000311028 | EAN76659 | 238 | *L.donovani* | ENSP00000300291 | XP_003865434.1 |
| 101 | *T.brucei* | ENSP00000323856 | EAN76659 | 239 | *L.donovani* | ENSP00000335321 | XP_003865434.1 |
| 102 | *Smansoni* | ENSP00000244601 | Smp_035980__mRNA | 240 | *L.donovani* | ENSP00000352336 | XP_003862574.1 |
| 103 | *Smansoni* | ENSP00000348924 | Smp_035980__mRNA | 241 | *L.donovani* | ENSP00000366179 | XP_003862230.1 |
| 104 | *Smansoni* | ENSP00000228434 | Smp_171460__mRNA | 242 | *L.donovani* | ENSP00000421592 | XP_003865434.1 |
| 105 | *Smansoni* | ENSP00000226730 | Smp_171460__mRNA | 243 | *L.donovani* | ENSP00000384144 | XP_003862230.1 |
| 106 | *Smansoni* | ENSP00000302648 | Smp_171460__mRNA | 244 | *L.donovani* | ENSP00000404042 | XP_003864767.1 |
| 107 | *Smansoni* | ENSP00000264690 | Smp_089670__mRNA | 245 | *L.donovani* | ENSP00000462972 | XP_003862223.1 |
| 108 | *Smansoni* | ENSP00000234071 | Smp_089670__mRNA | 246 | *L.braziliensis* | ENSP00000320866 | XP_001567285.2 |
| 109 | *Smansoni* | ENSP00000300289 | Smp_056760__mRNA | 247 | *L.braziliensis* | ENSP00000227378 | XP_001566161.1 |
| 110 | *Smansoni* | ENSP00000262262 | Smp_171460__mRNA | 248 | *L.braziliensis* | ENSP00000247461 | XP_001567285.2 |
| 111 | *Smansoni* | ENSP00000297185 | Smp_049550__mRNA | 249 | *L.braziliensis* | ENSP00000357998 | XP_001566161.1 |
| 112 | *Smansoni* | ENSP00000339007 | Smp_171460__mRNA | 250 | *G.lamblia* | ENSP00000357981-PF00112 | XP_001704582.1 |
| 113 | *Smansoni* | ENSP00000366962 | Smp_035980__mRNA | 251 | *G.lamblia* | ENSP00000357981-PF00112 | XP_001704877.1 |
| 114 | *Smansoni* | ENSP00000353074 | Smp_035980__mRNA | 252 | *G.lamblia* | ENSP00000327801 | XP_001708626.1 |
| 115 | *Smansoni* | ENSP00000369081 | Smp_056760__mRNA | 253 | *G.lamblia* | ENSP00000357981-PF00112 | XP_001708083.1 |
| 116 | *Smansoni* | ENSP00000365991 | Smp_049550__mRNA | 254 | *G.lamblia* | ENSP00000357981-PF00112 | XP_001706373.1 |
| 117 | *Smansoni* | ENSP00000340466 | Smp_018760__mRNA | 255 | *G.lamblia* | ENSP00000357981-PF00112 | XP_001709033.1 |
| 118 | *Smansoni* | ENSP00000327801 | Smp_056760__mRNA | 256 | *G.lamblia* | ENSP00000357981-PF08246 | XP_001704582.1 |
| 119 | *P.vivax* | ENSP00000223641 | PVX_003950 | 257 | *G.lamblia* | ENSP00000357981-PF028246 | XP_001709033.1 |
| 120 | *P.vivax* | ENSP00000215730 | PVX_003985 | 258 | *G.lamblia* | ENSP00000357981-PF08246 | XP_001704877.1 |
| 121 | *P.vivax* | ENSP00000249356 | PVX_099315 | 259 | *G.lamblia* | ENSP00000357981-PF08246 | XP_001706373.1 |
| 122 | *P.vivax* | ENSP00000227378 | PVX_099315 | 260 | *G.lamblia* | ENSP00000357981-PF08246 | XP_001708083.1 |
| 123 | *P.vivax* | ENSP00000054666 | PVX_003985 | 261 | *Chominis* | ENSP00000327801 | XP_667694.1-PF00085 |
| 124 | *P.vivax* | ENSP00000255194 | PVX_113995 | 262 | *Chominis* | ENSP00000327801 | XP_667694.1-PF13848 |
| 125 | *P.vivax* | ENSP00000259469 | PVX_003950 | 263 | *Chominis* | ENSP00000300289-PF00085 | XP_667694.1-PF13848 |
| 126 | *P.vivax* | ENSP00000262061 | PVX_113995 | 264 | *Chominis* | ENSP00000300289-PF13848 | XP_667694.1-PF13848 |
| 127 | *P.vivax* | ENSP00000249647 | PVX_003985 | 265 | *Chominis* | ENSP00000300289-PF13848 | XP_667694.1-PF00085 |
| 128 | *P.vivax* | ENSP00000261558 | PVX_113995 | 266 | *Chominis* | ENSP00000300289-PF00085 | XP_667694.1-PF00085 |
| 129 | *P.vivax* | ENSP00000304697 | PVX_119710 | 267 | *C.parvum* | ENSP00000264382-PF01055 | XP_627049.1-PF01055 |
| 130 | *P.vivax* | ENSP00000263354 | PVX_003985 | 268 | *C.parvum* | ENSP00000264382-PF00088 | XP_627049.1-PF01055 |
| 131 | *P.vivax* | ENSP00000294179 | PVX_003985 | 269 | *C.parvum* | ENSP00000264382-PF01055 | XP_627049.1-PF13802 |
| 132 | *P.vivax* | ENSP00000358126 | PVX_003985 | 270 | *C.parvum* | ENSP00000295688 | XP_628483.1 |
| 133 | *P.vivax* | ENSP00000287022 | PVX_117325 | 271 | *C.parvum* | ENSP00000300289-PF13848 | XP_625352.1-PF13848 |
| 134 | *P.vivax* | ENSP00000332887 | PVX_117325 | 272 | *C.parvum* | ENSP00000300289-PF13848 | XP_625352.1-PF00085 |
| 135 | *P.vivax* | ENSP00000367934 | PVX_117325 | 273 | *C.parvum* | ENSP00000325875-PF00183 | XP_628530.1-PF00183 |
| 136 | *P.vivax* | ENSP00000389103 | PVX_003950 | 274 | *C.parvum* | ENSP00000325875-PF02518 | XP_628530.1-PF00183 |
| 137 | *P.vivax* | ENSP00000354554 | PVX_117325 | 275 | *C.parvum* | ENSP00000325875-PF02518 | XP_628530.1-PF02518 |
| 138 | *P.vivax* | ENSP00000317159 | PVX_117325 | 276 | *C.parvum* | ENSP00000340466-PF13802 | XP_627049.1-PF01055 |

**Table S2. MassiveFold parameters used in the prediction of AF2-M models.**

| Parameter Name | Parameter Value | Complex no of complexes that run with these values |
| --- | --- | --- |
| model_preset | multimer | all |
| max_recycles | 3 | all |
| templates | true | all |
| max_res | 35000 | Human*-S.mansoni:* 104, 105, 106, 110, 112  *Human-L.mexicana: 212* |
| dropout | false | all |
| dropout_structure_module | false | all |
| dropout_recycling_below | 0 | all |
| min_score | 0 | all |
| max_score | 1 | all |
| db_preset | full_dbs | all |
| early_stop_tolerance | 0.5 | all |
| bfd_max_hits | 100000 | all |
| mgnify_max_hits | 501 | all |
| uniprot_max_hits | 50000 | all |
| uniref_max_hits | 10000 | all |
| MF_plots_top_n_predictions | 10 | all |
| MF_plots_chosen_plots | coverage,DM_plddt_PAE,CF_PAEs,score_distribution,recycles | all |

**Table S3. Percentage of cases with consistently low confidence scores (< 0.6) across all AlphaFold versions. Organisms are listed in descending order based on the frequency of these unsuccessful predictions.**

| Host-Parasite | Total Complex No | Percentage |
| --- | --- | --- |
| human-chominis | 6 | 100.00 |
| human-tbrucei | 15 | 93.33 |
| human-tcruzi | 54 | 88.89 |
| human-pknowlesi | 18 | 83.33 |
| human-pvivax | 18 | 83.33 |
| human-ldonovani | 12 | 83.33 |
| human-linfantum | 21 | 80.95 |
| human-cparvum | 10 | 80.00 |
| human-lmexicana | 35 | 74.28 |
| human-tspiralis | 7 | 71.42 |
| human-smansoni | 17 | 70.59 |
| human-tgondii | 25 | 48 |
| human-pfalciparum | 21 | 47.62 |
| human-glamblia | 11 | 45.45 |
| human-lbraziliensis | 4 | 25.00 |
